# Iron acquisition in the weevil-farming fungus *Penicillium herquei*: Implications of mineral elements in insect-fungus mutualistic association

**DOI:** 10.1101/2025.03.31.646393

**Authors:** Penglei Qiu, Xingzhong Liu, Dongsheng Wei

## Abstract

Mutualistic interactions between insects and fungi are pivotal in ecosystem dynamics, yet the underlying molecular mechanisms remain largely unexplored. This study investigates iron acquisition strategies of the weevil-farming *Penicillium herquei*, revealing the involvement of mineral elements in insect-fungus symbiosis. Comparative transcriptomics of weevil-farming strain (WFS) and soil free-living strain (SFS) revealed distinct transcriptional profiles, with 4,357 up-regulated genes in WFS. Enrichment analyses highlighted a significant up-regulation of genes linked to oxidoreductase activity, iron and heme binding, with a notable prevalence of Cytochrome P450 (CYP450). qRT-PCR confirmed differential expression of CYP450 and siderophore-related genes, indicating enhanced iron absorption in WFS. Comparative analysis of iron content further demonstrated significantly higher iron levels in WFS than in SFS and weevil host plant leaves, suggesting a nutritional adaptation for symbiotic lifestyle. These findings provide novel insights into the role of iron metabolism in insect-fungus mutualism, highlighting potential evolutionary mechanisms that bolster symbiotic fitness.

**IMPORTANCE:** Unraveling the complex interplay between insects and fungi is crucial for deciphering the intricate dynamics of ecosystems. In this study a notable up-regulation of genes associated with iron and heme binding, as well as a significant increase in iron content within WFS, suggesting a specialized adaptation for enhanced iron acquisition, potentially enabling the fungus to efficiently provide essential nutrients, including bioavailable iron, to weevil host. This research not only advances our understanding of the molecular mechanisms governing insect-fungus mutualism but also highlights the potential evolutionary mechanisms that bolster symbiotic fitness and contribute to the co-evolution of these interacting species.

## INTRODUCTION

Phytophagous insects frequently form mutualistic associations with fungi, facilitating fungal dispersal on plant substrates while benefiting from the fungi’s ability to provide accessible nutrients that promote insect growth (1‒4). These symbiotic relationships exhibit remarkable diversity, and their evolutionary persistence relies on intricate synergistic mechanisms. Such mutual interdependence, reinforced through recurrent coadaptation (5), has profoundly influenced Earth’s biodiversity by shaping ecological networks.

Fungus cultivation by insects, such as ants, termites, and ambrosia beetles, represents a sophisticated mutualistic symbiosis that shapes feeding niches and drives diversification among insect species (4). These insects function as adept practitioners of agricultural techniques, including the dispersal and seeding of fungal propagules, cultivation, and sustainable harvesting (6). Fungal crops provide diverse nutrients through specialized structures, such as fungal nodules in termite-farming fungi**^1^** and hyphal tip gongylidia in leaf-cutting ants (7). For example, attine ant fungal cultivars supply essential nutrients, including carbohydrates, vitamins, amino acids, and sterols, critical for insect survival (8). By cultivating fungi, these insects indirectly sustain themselves, as fungi convert lignocellulosic substrates into nitrogen-rich compounds (9‒12). Additionally, fungal symbionts may compensate for nutritional deficiencies in insect diets or provide chemical defenses against pathogens (7, 13, 14). Understanding the precise molecular mechanisms governing this mutual dependency remains a research priority with implications for symbiotic evolution and ecological adaptation.

The mutualistic association between the attelabid beetle *Euops chinensis* (Coleoptera: Attelabidae) and the fungus *Penicillium herquei* (Eurotiales: Aspergillaceae) represents a unique non-social insect-farming system (15). *E. chinensis*, prevalent in southern China, develops within curled fragments of *Reynoutria japonica* leaves, known as cradles, which serve as habitats for its eggs, larvae, and pupae. Beyond defense mechanisms and the vertical transmission of symbiotic *P. herquei* (14, 16), recent study demonstrated that fungal mycelia constitute a significant portion (62.6%-79.7%) of the diet of *E. chinensis* larvae, surpassing leaf materials, and providing essential nutrients such as ergosterol, amino acids, and B vitamins (17).

Our preliminary transcriptomic comparison between the weevil-farming strain (WFS) and soil free-living strain (SFS) of *P. herquei* revealed significant differences in iron metabolism. Studies have shown that insects feeding on nutritionally imbalanced diets, such as leaves and stems, have evolved mutualistic associations with fungi to acquire essential nutrients (1, 4, 7, 18). Iron, a vital microelement for both plants and insects (19, 20), is primarily found in plants as non-heme iron (e.g., ferritin, iron phosphate), which has low bioavailability (21, 22). In contrast, fungi, particularly lignocellulose-degrading genera, actively chelate environmental iron by secreting siderophores and accumulating it in their mycelia (23, 24). Thus, symbiotic *P. herquei* may facilitate iron acquisition by breaking down iron-bound compounds in *R. japonica* leaves and absorbing iron through siderophore secretion. This iron-enriched fungal biomass likely provides essential elements for the host weevil through a specialized iron acquisition strategy. This strategy was studied in details via deep analysis of comparative transcriptome and iron quantification in this study.

## MATERIALS AND METHODS

### Strain and plant materials

The weevil-farming strain Ph15277 (WFS) of *P. herquei* was isolated from the mycangium of female *E. chinensis*, while the soil free-living strain Ph5687 (SFS) was isolated from the surrounding environment. Both strains are preserved in our laboratory at Nankai University (Tianjin, China). Leaves of *R. japonica* were collected from Jiangxi Province, China.

### RNA extraction, cDNA library construction, and transcriptome sequencing

The fungal strains WFS and SFS were cultivated on potato dextrose agar at 26°C for seven days. Following cultivation, mycelia were collected (n = 3 for each strain), immediately frozen, and stored at -80°C for subsequent RNA-seq analysis. Total RNA was extracted from the frozen samples using the TRIzol method (Invitrogen, CA, USA). RNA degradation and contamination were assessed using 1% agarose gels. The concentration and purity of the RNA were measured with a NanoDrop spectrophotometer (Thermo Scientific, DE, USA), and RNA integrity was evaluated using an Agilent 2100 Bioanalyzer (Agilent Technologies, USA). A total of 1.5 μg RNA per sample was used for RNA sample preparation. Sequencing libraries were generated using the NEBNext Ultra™ RNA Library Prep Kit for Illumina (NEB, USA) according to the instructions of manufacturer. The library preparations were sequenced on an Illumina NovaSeq 6000 platform by Beijing Allwegene Technology Company Limited (Beijing, China), producing paired-end 150 bp reads.

### Mapping analysis and quantification of gene expression levels

Clean reads were obtained by filtering out adapter-containing sequences, poly-N sequences, and low-quality reads from the raw data. The quality metrics, including Q20, Q30, GC content, and sequence duplication level, were calculated for the clean data. All downstream analyses were based on this high-quality clean data. The clean reads were mapped to the reference genome sequence of the weevil-farming strain (Ph15277) using STAR. Only reads with a perfect match or a single mismatch were analyzed and annotated based on the reference genome. Read counts for each gene were obtained using HTSeq v0.5.4 p3, and gene expression levels were estimated as fragments per kilobase of transcript per million fragments mapped (FPKM) (25).

### Analysis of differentially expressed genes (DEGs)

Differential expression analysis between the two fungal groups was conducted using the DESeq R package (1.10.1). The resulting P-values were adjusted using the Benjamini and Hochberg method to control the false discovery rate. Genes with an adjusted P-value < 0.05 identified by DESeq were classified as differentially expressed. Gene Ontology (GO) enrichment analysis of the differentially expressed genes (DEGs) was performed using the GOseq R package based on the Wallenius non-central hypergeometric distribution (26). The KOBAS software was utilized to evaluate the statistical enrichment of DEGs in KEGG pathways (27).

### qRT-PCR analysis

The fungal strains WFS and SFS were maintained on potato dextrose agar at 26°C for seven days. Mycelia were collected post-cultivation, immediately frozen, and stored at -80°C for subsequent RNA-seq analysis. Total RNA was extracted from these frozen samples using TRIzol reagent (Thermo Fisher Scientific, USA). The cDNA was synthesized from 1 μg of total RNA using the TransScript All-in-One SuperMix reagent kit (TransGen, China). For quantitative PCR (qPCR), 0.2 μL of cDNA was used as the template, employing PerfectStart Green qPCR SuperMix (TransGen, China) with the CFX Connect real-time system (Singapore), according to the manufacturer’s guidelines. The actin gene served as an internal control for normalizing the relative expression levels of each gene. Final calculations were performed using the threshold cycle (2^−ΔΔCT^) method (28).

### Determination of iron content

Iron content was measured by digesting 0.18 g of dry-weight plant leaves from *R. japonica* and 0.1 g of dry-weight mycelia from WFS and SFS in concentrated HNO_3_ at 65°C for 45 minutes (29). The resultant samples were then heated at 92°C to remove residual HNO_3_. Following this, 5 mL of deionized water was added to the mycelia samples, and 2.5 mL of deionized water was added to the plant samples. The solutions were filtered through a 0.22 μm membrane prior to inductively coupled plasma (ICP) detection.

## RESULTS

### RNA sequencing and transcriptomic assembly

After filtering raw data (NCBI SRA Bioproject ID: PRJNA1225116), we obtained 43.47 Gb of clean data from six cDNA libraries, which included three from the weevil-farming strain (WFS) and three from the soil free-living strain (SFS). The average yield per sample was 6.96 Gb for WFS and 7.52 Gb for SFS, with total reads averaging 46,438,696 and 50,172,612, respectively. High-quality reads exhibited an average GC content of 51.73%, with Q20 and Q30 scores exceeding 98% and 94%, respectively (Table 1), ensuring data suitability for further analysis. Read alignment rates to the WFS reference genome (Ph15277) ranged from 81.89% to 95.27%. Principal component analysis (PCA) confirmed strong clustering of biological replicates (Fig. 1A), with Pearson correlation coefficients exceeding 0.92 (Fig. 1B), indicating high-quality and consistent RNA-seq data.

**FIG 1.**
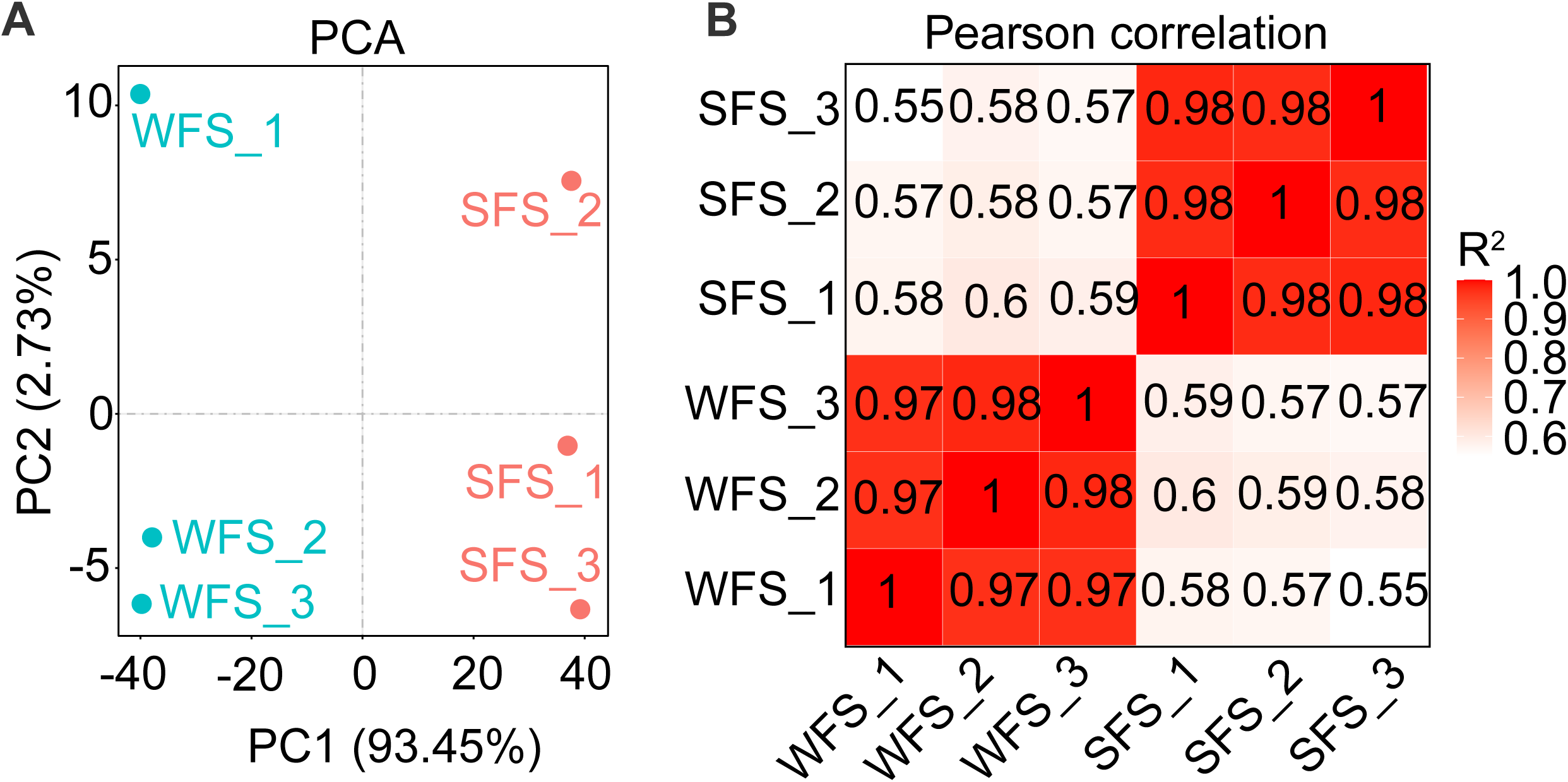
Evaluation of correlations across weevil-farming strain (WFS) and soil-free living strain (SFS) of *Penicillium herquei*. (**A‒B**) Principal component analysis (**A**) and Pearson correlation coefficients (**B**) of WFS and SFS samples.

**TABLE 1.** RNA-Seq statistics of weevil-farming strain (WFS) and soil-free living strain (SFS) of *Penicillium herquei*.

| Sample | Clean reads | Clean bases (G) | % Q20 | % Q30 | % GC | % Mapped |
| --- | --- | --- | --- | --- | --- | --- |
| WFS_1 | 48356356 | 7.25 | 98.08 | 94.36 | 51.78 | 95.06 |
| WFS_2 | 46199062 | 6.93 | 98.11 | 94.47 | 51.62 | 95.2 |
| WFS_3 | 44760670 | 6.71 | 98.13 | 94.55 | 51.59 | 95.27 |
| SFS_1 | 51106902 | 7.67 | 98.01 | 94.30 | 51.66 | 81.89 |
| SFS_2 | 46458634 | 6.97 | 98.12 | 94.59 | 51.98 | 83.06 |
| SFS_3 | 52952302 | 7.94 | 98.08 | 94.49 | 51.76 | 81.99 |

### Identification of differentially expressed genes (DEGs)

To explore the molecular basis of mutualistic symbiosis, we conducted a comparative transcriptome analysis between WFS and SFS. The DEGs were identified based on a fold change (fc) greater than 2 and a p-value less than < 0.05. Among the 12,987 annotated genes, 4,357 were significantly up-regulated and 3,258 were down-regulated in WFS compared to SFS (Fig. 2A; Table S1). Additionally, we identified 9,644 co-expressed genes, with WFS displaying a higher number of strain-specific expressed genes than SFS (Fig. 2B; Table S2). A heatmap of all DEGs highlighted distinct transcriptional profiles between WFS and SFS (Fig. 2C).

**FIG 2.**
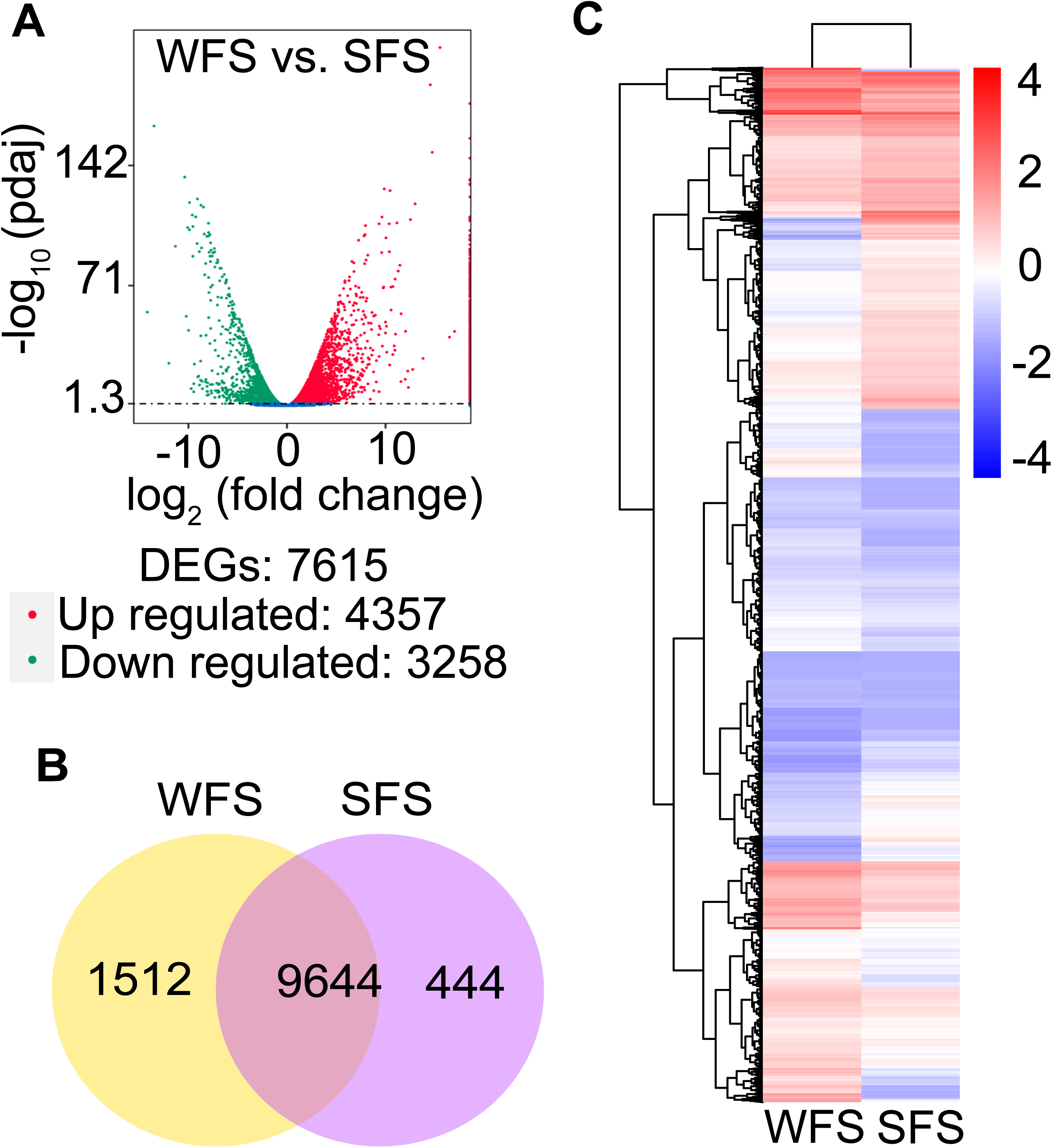
Differential gene expression analysis of weevil-farming strain (WFS) and soil free-living strain (SFS) of *Penicillium herquei*. (**A‒C**) The volcano map (**A**), Venn diagram (**B**) and heatmap (**C**) of differentially expressed genes for WFS and SFS samples.

### GO and KEGG enrichment analysis of DEGs

Gene Ontology (GO) annotation of DEGs assigned 6,177 unigenes to 3,474 GO terms (Table S3). The DEGs were predominantly associated with “molecular function”, followed by “biological processes” and “cellular components”. Notably, a significant number of up-regulated DEGs were enriched in GO terms related to oxidoreductase activity, iron binding, and heme binding, as well as oxidation-reduction processes (Fig. 3A and B). In contrast, down-regulated DEGs were mainly involved in metabolic and biosynthetic processes (Fig. S1).

**FIG 3.**
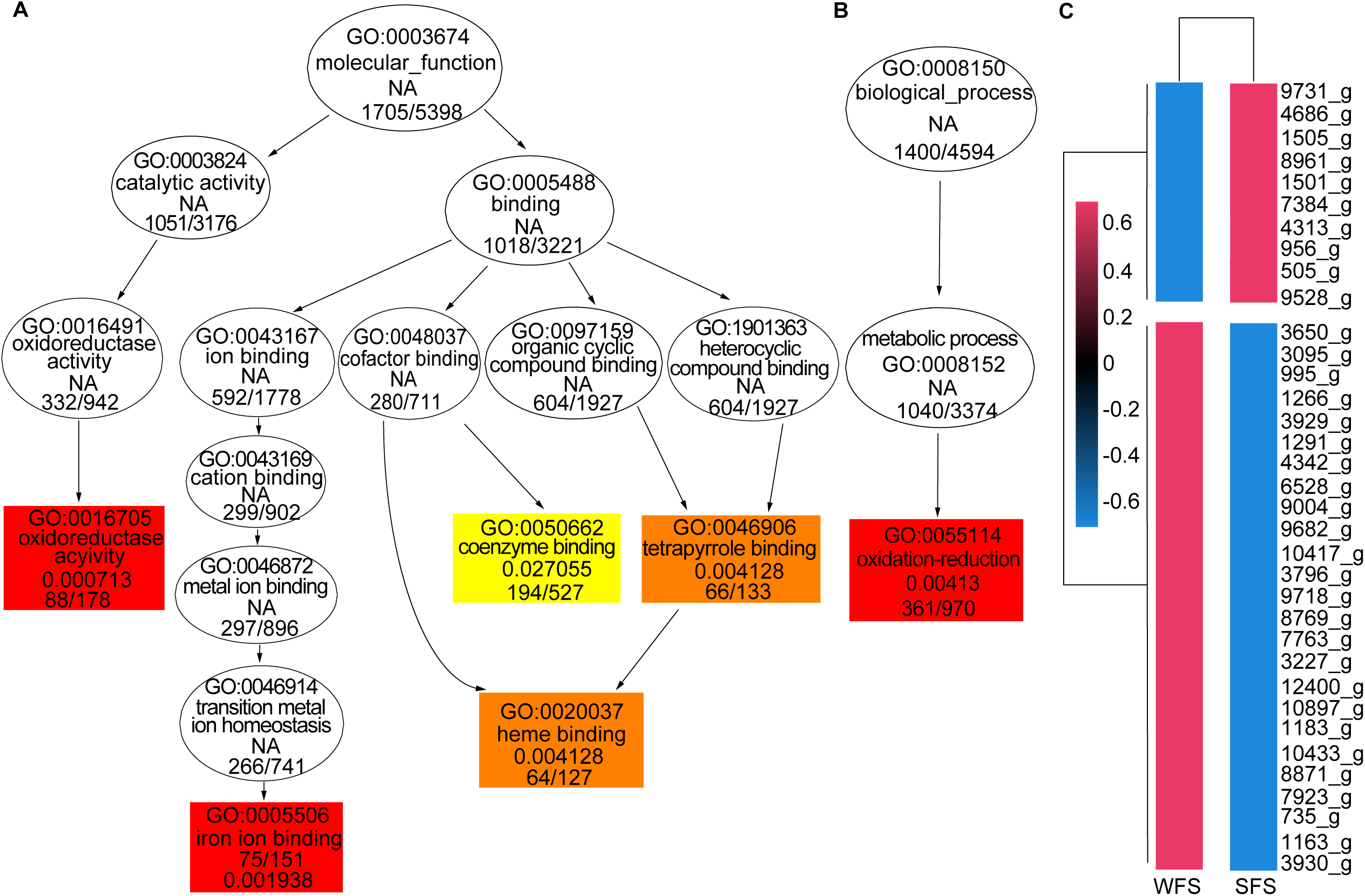
Enrichment of the differentially expressed genes (DEGs) for weevil-farming strain (WFS) and soil free-living strain (SFS) of *Penicillum herquei*. (**A‒B**) GO enrichment analysis of differentially up-regulated genes in molecular functions (**A**) and biological processes (**B**). (**C**) Cluster heatmap analysis of differentially expressed CYP450 genes for WFS and SFS.

Notably, most up-regulated genes related to iron (75 genes) and heme (64 genes) binding encoded Cytochrome P450 (CYP450) proteins (Table S3), with over 70% of CYP450 genes significantly up-regulated (Fig. 3C). Additionally, the KEGG pathway analysis revealed enrichment of up-regulated DEGs in fatty acid and steroid biosynthesis, alongside a variety of metabolic pathways involving amino acids, saccharides, and fatty acids (Fig. S2A). Conversely, down-regulated DEGs were predominantly associated with the ribosome pathway (Fig. S2B).

### qRT-PCR validation of CYP450 and iron-related gene expression

To validate RNA-seq results, we randomly selected four CYP450 genes for qRT-PCR analysis, which confirmed expression trends consistent with RNA-seq data (Fig. 4A‒D). To further compare iron absorption between WFS and SFS, we analyzed the expression of four siderophore-related genes. These genes were significantly up-regulated in WFS (Fig. 4E‒H), indicating enhanced iron uptake capacity.

**FIG 4.**
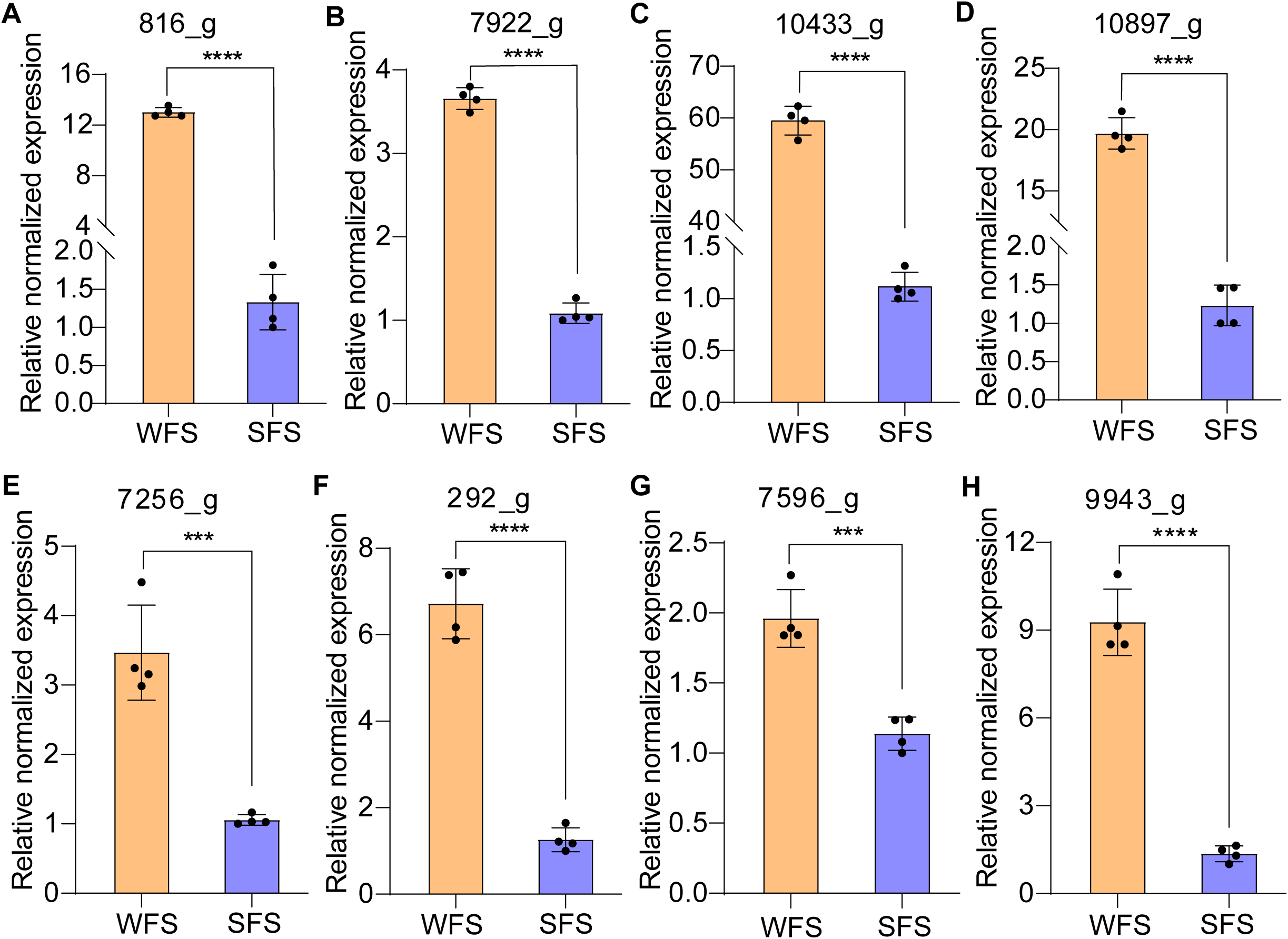
Quantitative analysis of expression for iron-binding and siderophore related genes. **(A‒D)** RT-qPCR for catalase (**A**), monoxygenase (**B**), benzoate 4-monooxygenase cytochrome p450 (**C**) and Cytochrome P450 (**D**). **(E‒H)** RT-qPCR for siderophore synthetase (**E**), siderophore transporter (**F**), siderochrome-iron transporter (**G**) and siderophore esterase (**H**).

### Comparative analysis of iron content in WFS, SFS, and plant leaves

Iron content analysis confirmed significantly higher iron levels in WFS compared to SFS (Fig. 5). Given the dietary preference of weevils for fungal mycelia over plant leaves (17), we also measured iron concentrations in *R. japonica* leaves. Results revealed lower iron levels in leaves than in fungal mycelia, highlighting the nutritional advantage of mycelia for iron intake. Additionally, weevil-farming fungal mycelia exhibited higher iron content than soil free-living strain, suggesting an evolutionary adaptation in WFS to enhance iron accumulation for its symbiotic lifestyle. This finding underscores the critical role of iron in this mutualistic interaction.

**FIG 5.**
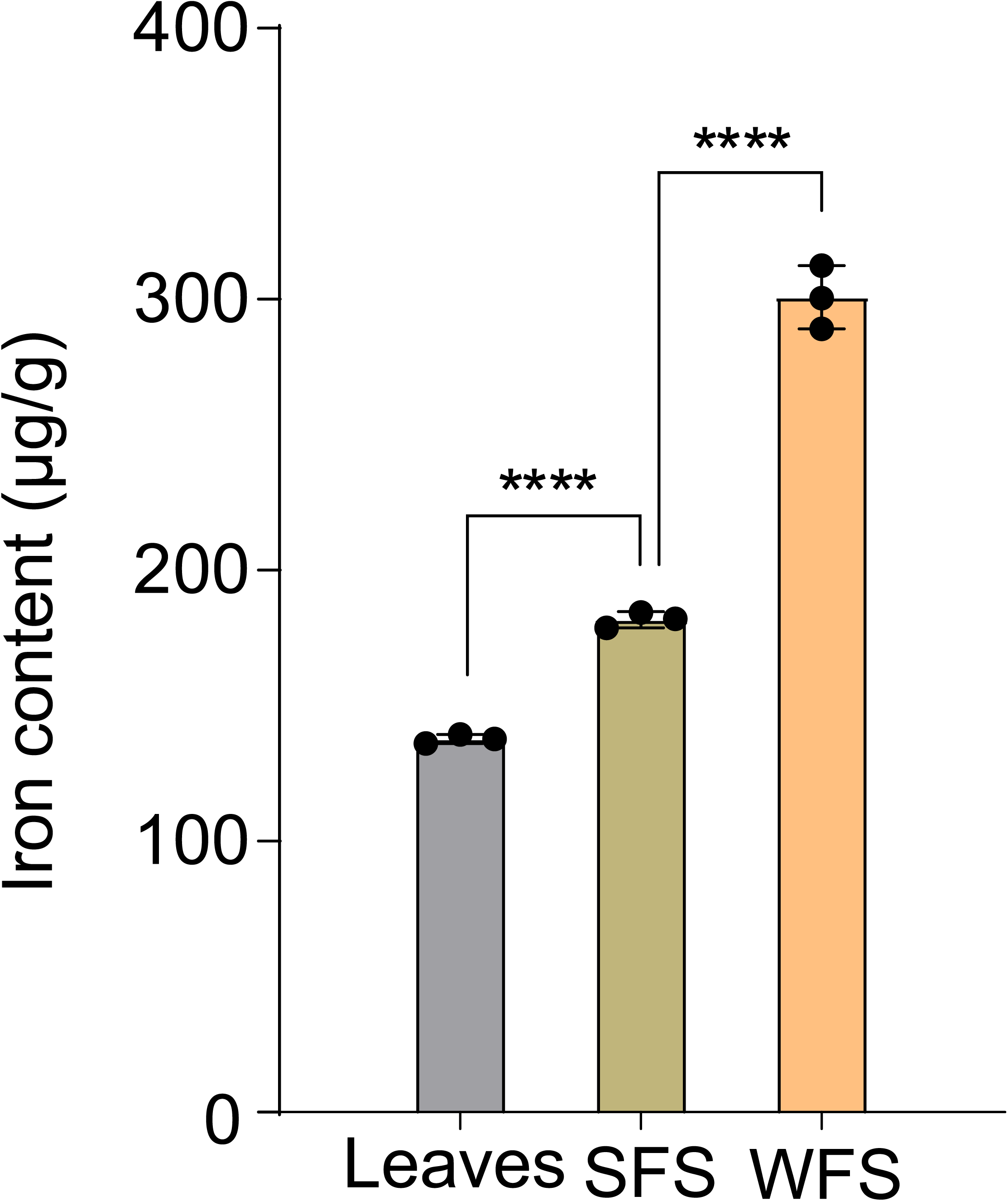
Determination of iron content in leaves of *Reynoutria japonica*, soil free-living strain (SFS) and weevil-farming strain (WFS) of *Penicillum herquei*.

## DISCUSSION

In well-characterized termite and attine ant fungiculture systems, specialized fungal structures, such as nodules and gongylidia, facilitate the delivery of essential amino acids and sterols crucial for larval development (1, 7). Comparisons of the nutrient profiles of the weevil-farming fungus and leaf rolls confirm that *P. herquei* plays a vital nutritional role, supplying the weevil with key compounds like ergosterol, amino acids, and B vitamins (17). Our findings further highlight the critical role of mineral elements, particularly iron, in the mutualistic symbiosis between insect and fungus, revealing an often-overlooked dimension of this relationship.

Iron (Fe) is a limiting micronutrient for many herbivores, directly influencing their growth and development (30). While insect herbivores typically obtain iron by consuming plant material, fungus-farming insects rely on fungal tissue to fulfill their micronutrient needs, including essential trace elements. In biological systems, iron is primarily bound to proteins to prevent precipitation and mitigate potentially harmful chemical reactions (31). Within *P. herquei*, Cytochrome P450 enzymes, a superfamily of heme-containing proteins, play a crucial role in both anabolic and catabolic pathways, underscoring their significance in biochemical processes (32). The enhanced capacity of Cytochrome P450 for iron or heme binding in *P. herquei* not only highlights its metabolic versatility but also provides substantial nutritional benefits to the weevil. Our findings, which link the expression of iron absorption and binding genes with iron quantification in fungal mycelia and plant leaves, confirm that insects can obtain sufficient iron by feeding on fungal mycelia, which contain significantly higher iron levels than the leaves of *R. japonica*.

Weevils benefit from the high bioavailability of iron in *P. herquei*, which likely enhances their growth and survival. Additionally, the mycelia contain elevated iron levels and release more soluble iron in the hypoxic and acidic conditions of the weevil midgut, facilitating iron assimilation during digestion (33, 34). However, increased iron availability could also impose stress on midgut microbiota, potentially inhibiting microbial proliferation (35). Interestingly, our unpublished data suggest that *P. herquei* has evolved to secrete Pc protein, which binds to and inhibits the activity of bacterial extracellular autolysin containing a beta-N-acetylglucosaminidase (GL) domain, thereby selectively enriching beneficial gut microbiota through mitigating iron and reactive oxygen species (ROS) stress.

The elevated iron content in the weevil-farming fungus supports the hypothesis that Pc protein serves as a key regulator of weevil gut microbiota, influencing microbial balance and overall digestive health. This specialized adaptation of enhanced iron absorption in *P. herquei* may represent a critical evolutionary strategy that optimizes nutrient exchange and mutualistic benefits in the weevil-fungus symbiosis. These findings contribute to a broader understanding of the metabolic and ecological mechanisms governing insect-fungal mutualisms, highlighting iron as a pivotal element in the co-evolution of these interacting species.

### Conclusions

This study highlights the crucial role of iron acquisition in the mutualistic symbiosis between *P. herquei* and weevil insects. While previous research has primarily focused on the provision of amino acids, sterols, and vitamins by fungal symbionts, our findings reveal that *P. herquei* also serves as a key source of bioavailable iron, surpassing the iron content found in host plant leaves. The up-regulation of Cytochrome P450 genes and siderophore-related genes in the weevil-farming strain suggests a specialized adaptation to enhance iron acquisition, benefiting the nutritional needs of its insect partner.

## ACKNOWLEDGMENTS

This work was supported by a grant from the Startup Fund from the Nankai University to X.L. (030/C029215002). We thank Luwen Yan for providing the leaf material of *Reynoutria japonica*.

## AUTHOR AFFILIATIONS

^1^Key Laboratory of Molecular Microbiology and Technology of the Ministry of Education, Department of Microbiology, College of Life Science, Nankai University, Tianjin 300071, China.

## AUTHOR CONTRIBUTIONS

Xingzhong Liu, Conceived and supervised the project | Penglei Qiu, Performed experiments and analyzed the data | Penglei Qiu, Xingzhong Liu and Dongsheng Wei, Co-wrote the manuscript, incorporating input from all authors.

## DATA AVAILABILITY

The raw data for transcriptome sequencing have been deposited in NCBI database under the accession number PRJNA1225116. This study did not generate new unique reagents. All the materials are available on request to the lead contact, Xingzhong Liu.

